# Methods for combining multiple correlated biomarkers with application to the study of low-grade inflammation and muscle mass in senior horses

**DOI:** 10.1101/2019.12.18.881425

**Authors:** Olga A. Vsevolozhskaya, Alisa C. Herbst, Amanda A. Adams, Cailey Burns, Bertsie Cantu, Virginia D. Barker, Dmitri V. Zaykin

## Abstract

The simplest analysis of biomarker data is based on a series of single biomarker hypothesis tests, followed by correction for multiple testing. However, it is intuitively plausible that a joint analysis of multiple biomarkers will have higher statistical power and promise improved discrimination over tests based on single markers. In this article, we study analytical properties of the approach for joint analysis of correlated summary statistics based on the test for quadratic forms (TQ). Based on the derivation of the TQ-distribution, we proposed a scale-location approximation of the TQ statistic, which we call approximate TQ. We show that the approximate TQ has very similar power to the traditional TQ test under varying correlation structures among biomarkers. Our application of both the TQ and the approximate TQ test to data on biomarkers for inflamm-aging – an age-related low-grade chronic inflammation – reveals an association between the percentage of IFNγ positive lymphocytes and overall muscle condition in senior horses.

## Introduction

The role of biomarkers in biomedical research has been steadily increasing over the recent years. For example, a key aspect of research in animal or human genetics is association of single nucleotide polymorphisms (SNP biomarkers) with complex traits; in large-scale gene expression analyses, one’s goal is to identify differentially expressed gene biomarkers; in cancer screening, increasing efforts are put into identification of “cancer biomarkers” – specific proteins released by tumor into blood; in magnetic resonance imaging (MRI), MRI lesions are studied as potential predictors of further disease progression, etc. In modern large-scale studies, the number of biomarkers collected can greatly exceed the number of samples. Therefore, each biomarker is typically assessed individually for its potential association with disease status. At the same time, the notion that combinations of multiple biomarkers may provide higher diagnosis accuracy is becoming widely accepted.

A number of statistical methods exists that can combine biomarker-specific summary statistics (e.g., Z-scores or P-values) and assess an association between multiple biomarkers and a trait. The Fisher combined probability test is perhaps one of the most popular ones (Fisher, 1992). Despite its wide popularity, the Fisher test relies on the assumption of independence among summary scores, while measured levels of many of the biomarkers are expected to be correlated with each other. For example, minor allele counts are expected to be correlated among SNPs and this correlation is captured by linkage disequilibrium (LD); gene expression levels are expected to be correlated in subsets of informative genes, etc. Therefore, when biomarker-specific association statistics are being combined, the correlation among biomarkers needs to be accounted for, because it induces correlation on the summary statistics.

Multiple correlated statistics can be combined using the test for quadratic forms (TQ). Under TQ, P-values for *p* individual-level biomarkers are first transformed into Z-scores, *P_i_* → *Zp_i_*, *i* = 1,…, *p*, and then combined as 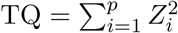. In this paper, we explore theoretical properties of the TQ test by deriving its distribution under the alternative hypothesis. Based on this derivation, we provide an approximation to the TQ test that is just as powerful as the original test. Our approximation will be very useful to researchers who have summary statistics for individual biomarkers available but no pair-wise correlation values. Finally, we showcase both the TQ and the approximate TQ test by applying them to data on biomarkers for inflamm-aging – an age-associated, low-grade and chronic inflammation (Franceschi et al., 2000), – and muscle mass in senior horses.

## Methods and Materials

### The relationship between correlation among biomarkers and estimated biomarker effects

Let y′ = (*y*_1_,…, *y_n_*) be an *n* × 1 vector of trait values and **X** be a *n* × *p* matrix of measured biomarkers. Suppose that **R**: *p* × *p* is the correlation matrix among biomarkers with *R_ij_* = Cor(*X_i_, X_j_*), where *X_i_* and *X_j_* are *i*th and *j*th columns of **X** respectively. A vector of Z-scores (estimated effect sizes divided by their standard errors) obtained by marginally testing each biomarker one at a time can be expressed in terms of the expected correlations between predictor *X_j_* and the outcome y as:

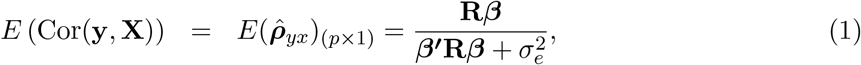

where *β*: *p*× 1 is a vector of biomarker effects and 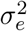 is defined by the linear model 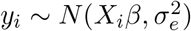. If **R** = **I**_*p*_ is an identity matrix (representing an unlikely scenario of uncorrelated biomarkers), expected univariate correlations become proportional to the vector of biomarker effects, 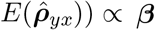. That is, a zero coefficient *β_i_* would imply zero expected correlation between biomarker *X_i_* and the outcome. When correlation among biomarkers is present, expected univariate correlations become proportional to weighted sums of all *β*, 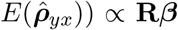, with weights given by the columns *R_j_*’s. Thus, on the one hand, a biomarker with no effect on the outcome (*β_i_* = 0) can still be correlated with it (*ρ_yi_* ≠ 0) due to the correlation with causal biomarkers. If a biomarker represents a SNPs, such non-associated SNPs are typically referred to as “proxy SNPs” in literature (Spencer et al., 2009). On the other hand, marginally testing a truly causal biomarkers (*β_i_* ≠ 0) may result in an estimated negligible effect (*ρ_yi_* ≈ 0) if *R_i_β* is close to zero.

Construction of multiple-biomarker association P-value requires that test statistics in Eq. (1) be asymptotically distributed as multivariate normal random variables with a known correlation matrix. When none of the biomarkers are associated with a trait, the common correlation matrix among association statistics is a function of correlation among biomarkers. Specifically, if univariate regression models do not include covariates, then correlation among biomarker-level statistics will be equal to the correlation matrix among them, Cor(*R_yi_*, *R_yj_*) → Cor(*X_i_*,*X_j_*) = *R_ij_*. With additional covariates that may be correlated with biomarkers (e.g., age, sex), Cor(*R_yi_*, *R_y_j*) will be a Schur complement of the submatrix **X** of the matrix of all predictor variables (Conneely and Boehnke, 2007). However, if some of the biomarkers are associated (or even if all of them are only proxies for the causal biomarkers), the problem of finding the common correlation matrix among association statistics becomes more challenging.

### The relationship between correlation among biomarkers and correlation among test statistics under the alternative hypothesis

We relate the problem of estimating a common correlation matrix among association statistics to the problem of correlation among correlations, which resurfaces throughout the history of statistics in diverse applications, starting with the 1898 paper by Pearson and Filon (Pearson et al., 1898). To derive the variance-covariance matrix of the test statistics, we assume that our sample consists of *n* vectors from a (*p* + 1)-variate multinormal distribution, (*y_i_*, *x*_1*i*_,…, *x_pi_*)′, with covariance matrix that has the following structure:

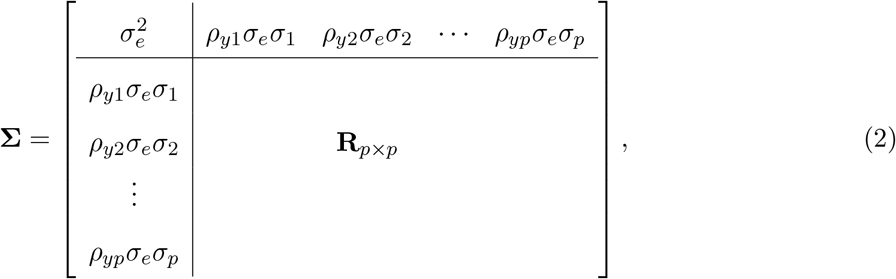

where 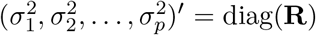 and (*ρ*_*y*1_, *ρ_y2_*,…, *ρ_yp_*)′ = ***ρ_yx_*** are pair-wise correlations between the outcome and *p* biomarkers. Since summary statistics can be expressed in terms of the expected correlations (Eq. (1)), to find a covariance matrix among statistics one needs to find the covariance matrix of the correlation estimates 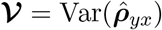.

Suppose we ignore the structure in Eq. (2) and just calculate the sample covariance matrix **V**. Diagonal elements of sample covariance matrix, *v_ii_*, are the maximum likelihood estimates of 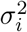 and off-diagonal elements, *v_ij_*, are the maximum likelihood estimates of *ρ_ij_σ_i_σ_j_*. Let 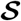 be the covariance matrix of the vectorized sample covariance matrix, 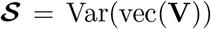 (Elston, 1975). The properties of a multivariate normal distribution ensures that if *s_ij_* is an estimate of the (*i,j*)th element, *c_ij_*, of the covariance matrix, then 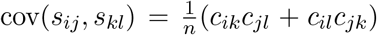 (e.g., Kendall et al. (1946)). Using this property, we can estimate the elements of 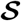 as follows:

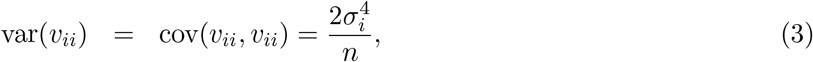

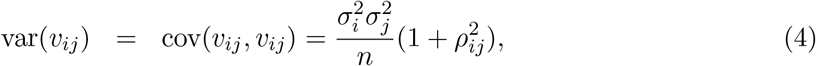

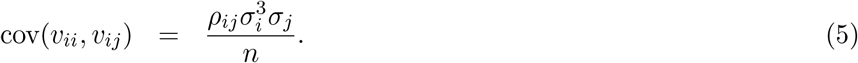

Finally, we can estimate variances and covariances of correlations using the Delta method as 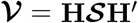, where the (*i,j*)’th element of **H** is the derivative of the *i*th estimated correlation with respect to the *j*th element of vec(**V**), evaluated at the parameter values. For example, the first row of **H** will contain partial derivatives of 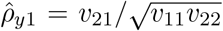 with respect to all *v_ij_*. Note that only 3 of these partial derivatives will have non-zero values: (1) 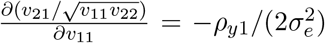, (2) 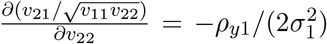, and (3) 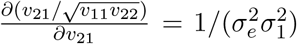. Similarly, the last row of **H** will contain partial derivatives of 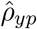 with respect to all *v_ij_*. Once again, only three of these partial derivatives will have non-zero values. Therefore, the matrix **H** can be constructed as:

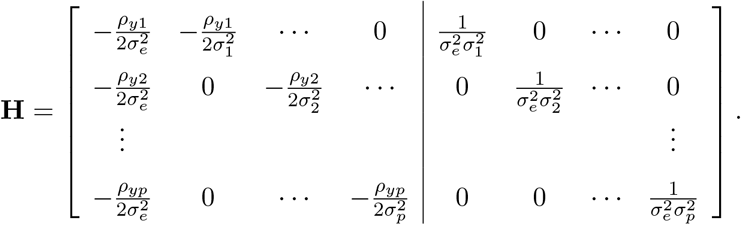

After multiplying matrices and simplification, diagonal and off-diagonal elements of 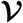 are found as:

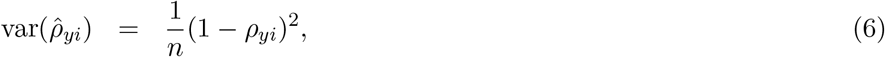

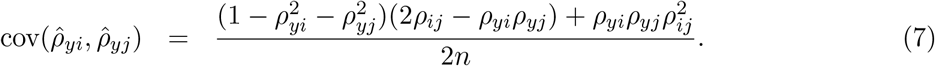

In our original notation, *ρ_yi_* = *b_i_* is the true value of the standardized regression coefficient, and *ρ_ij_* = *R_ij_* is the correlation matrix among biomarkers. Thus,

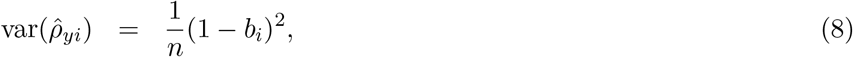

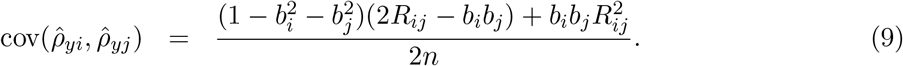

Further, a good approximation to covariances and correlations can be obtained via:

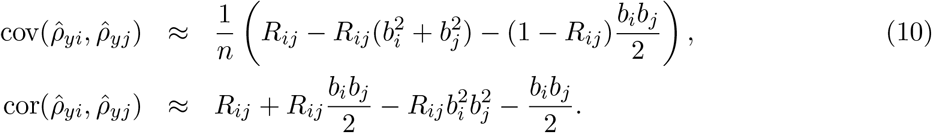

### Multiple Biomarker test

Multiple biomarker-based tests aggregate evidence for an association across all *p* biomarkers by computing a weighted sum of single-biomarker Z-scores (burden tests) or a weighted sum of squared single-biomarker Z-scores (e.g., overdispersion tests). Let 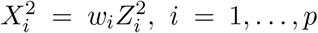, denote a weighted squared Z-score. Based on the previous section, we know that 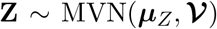 with ***μ***_*z*_ = (*ρ*_*y*1_,…, *ρ_yp_*)′ and thus 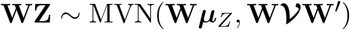, where 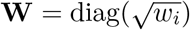. The multiple biomarker-based statistic is defined as 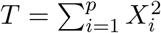. The values of 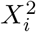’s are correlated but following standard theory, the distribution of their sums can be represented as the distribution of the weighted sum of independent one degree of freedom chi-squares:

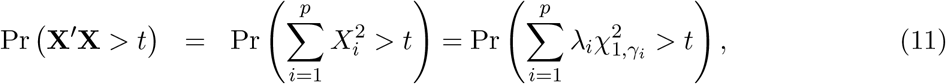

where λ_*i*_ is the ith eigenvalue of 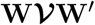, and γ*_i_* is a non-centrality parameter defined as 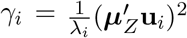, where **u***_i_* is the ith orthonormal eigenvector of 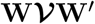. The P-value for the statistic *T* = **X**′**X**, calculate under the global null hypothesis of no association, is obtained by setting **μ***_Z_* to zero, substituting 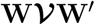 by **WRW**′ and then calculating the tail probability at the observed value *T* = *t*.

### Power of the multiple Biomarker test

To examine statistical power, in contrast to the null model (that is, **Z** ~ MVN(**0**, **R**)) we need to consider the case were causal biomarker have non-zero effects (that is 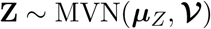). For now, let’s assume that 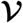 is equicorrelated with common correlation, *ν* > 0, for all pairs of statistics.

Then, 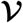 will have only two distinct eigenvalues:

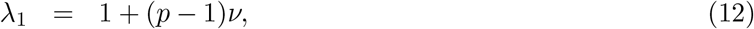

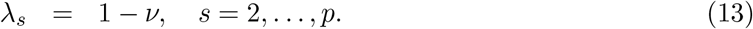

The orthonormal eigenvectors can be obtained by normalizing the orthogonal basis:

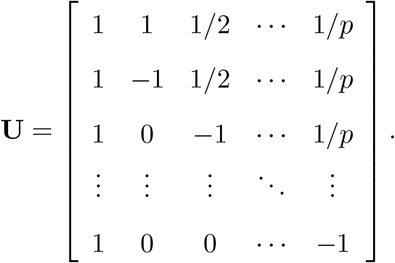

The corresponding non-centralities can be derived as follows:

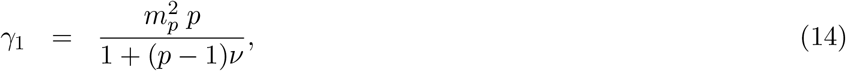

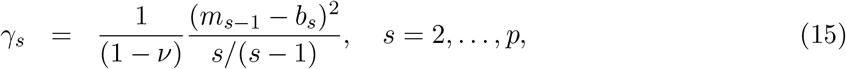

where 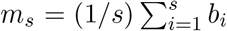. Consider the behavior of non-centralities as the number of biomarkers to be jointly tested goes to infinity. On the one hand, the first non-centrality increases with *p*:

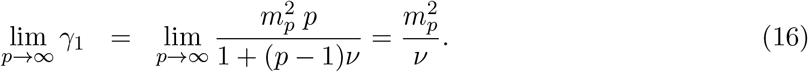

On the other hand, the sum of the remaining non-centralities, 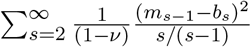, does not di verge, which is easy to show by the ratio test. Therefore, under the null hypothesis, the distribution of the quadratic form **X**′**X** can be well approximated by the location-scale transformation of the one degree of freedom chi-squared random variable:

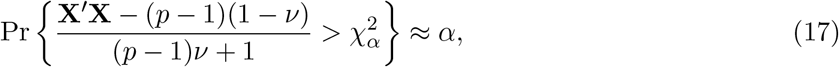

where 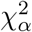 is the 1 – *α* quantile of the one degree of freedom chi-square distribution.

## Results

### Simulation study

We conducted simulation experiments to study statistical power of the proposed approximation to the TQ test (Eq. (17)) in relation to the traditional TQ test statistic (Eq. (11)). Tables 1 and 2 summarize the results.

**Table 1:** Type I error of TQ and approximate TQ methods. Simulations were performed under complete null hypothesis (Pr(*H*_0_) = 100%) but with different correlation structures among biomarkers. The value *ν* indicates the range of values of the off-diagonal elements of the correlation matrix **R** among biomarkers. Specifically, *ν* = 0.5 represents the case of equi-correlated structure with all off-diagonal elements equal to 0.5; *ν* ∈ (−0.25, 0.75) represents a general correlation matrix with the range of the off-diagonal values varying between –0.25 and 0.75; *ν* ∈ (–0.70,0.90) represents a general correlation matrix with a higher level of heterogeneity among its element. Different values of *p* indicate the number of biomarkers that were combined.

| $\Pr(H_0) = 100\%, \nu = 0.5$ | | | | |
| --- | --- | --- | --- | --- |
| | $p = 50$ | $p = 30$ | $p = 20$ | $p = 10$ |
| Empiric. TQ | 0.047 | 0.051 | 0.047 | 0.053 |
| Approx. TQ | 0.047 | 0.054 | 0.049 | 0.051 |
| $\Pr(H_0) = 100\%, \nu \in (-0.25, 0.75)$ | | | | |
| | $p = 50$ | $p = 30$ | $p = 20$ | $p = 10$ |
| Empiric. TQ | 0.044 | 0.052 | 0.045 | 0.045 |
| Approx. TQ | 0.046 | 0.052 | 0.044 | 0.046 |
| $\Pr(H_0) = 100\%, \nu \in (-0.70, 0.90)$ | | | | |
| | $p = 50$ | $p = 30$ | $p = 20$ | $p = 10$ |
| Empiric. TQ | 0.053 | 0.048 | 0.055 | 0.048 |
| Approx. TQ | 0.052 | 0.049 | 0.055 | 0.058 |

**Table 2:** Power comparison of TQ and approximate TQ methods. Simulations were performed under the assumption that 80% of the measured biomarkers were not associated with the trait (Pr(*H*_0_) = 80%). Effect sizes of biomarkers that were truly associated were drawn from a unifrom distribution within bounds that either corresponded to little effect size heterogeneity (i.e., −0.1, 0.1) or higher effect size heterogeneity (i.e., −0.2, 0.2). Correlation structure among biomarkers followed the same pattern as in Type I error simulations.

| $\Pr(H_0) = 80\%, b_i \sim \text{Unif}(-0.1, 0.1), \nu = 0.5$ | | | | |
| --- | --- | --- | --- | --- |
| | $p = 50$ | $p = 30$ | $p = 20$ | $p = 10$ |
| Empiric. TQ | 0.537 | 0.440 | 0.372 | 0.261 |
| Approx. TQ | 0.536 | 0.448 | 0.377 | 0.263 |
| $\Pr(H_0) = 80\%, b_i \sim \text{Unif}(-0.1, 0.1), \nu \in (-0.25, 0.75)$ | | | | |
| | $p = 50$ | $p = 30$ | $p = 20$ | $p = 10$ |
| Empiric. TQ | 0.594 | 0.492 | 0.395 | 0.254 |
| Approx. TQ | 0.602 | 0.485 | 0.392 | 0.267 |
| $\Pr(H_0) = 80\%, b_i \sim \text{Unif}(-0.2, 0.2), \nu \in (-0.25, 0.75)$ | | | | |
| | $p = 50$ | $p = 30$ | $p = 20$ | $p = 10$ |
| Empiric. TQ | 0.861 | 0.798 | 0.717 | 0.588 |
| Approx. TQ | 0.865 | 0.800 | 0.729 | 0.582 |
| $\Pr(H_0) = 80\%, b_i \sim \text{Unif}(-0.2, 0.2), \nu \in (-0.70, 0.90)$ | | | | |
| | $p = 50$ | $p = 30$ | $p = 20$ | $p = 10$ |
| Empiric. TQ | 0.879 | 0.820 | 0.757 | 0.616 |
| Approx. TQ | 0.876 | 0.819 | 0.755 | 0.609 |

Table 1 demonstrates type I simulation results. Simulations were performed under complete null hypothesis (i.e., Pr(*H*_0_) = 100%) but with different correlation structures **R** among biomarkers. The value v indicates the range of values of the off-diagonal elements of **R**. For example, *ν* = 0.5 represents the case of equi-correlated structure with all off-diagonal elements equal to 0.5, *ν* ∈ (–0.25,0.75) represents a general correlation matrix with the range of the off-diagonal values varying between −0.25 and 0.75, *ν* ∈ (–0.70, 0.90) represents a general correlation matrix with a higher level of heterogeneity among its element. These different general correlation matrices were constructed by adding rank-one matrix **U**′**U** to the equi-correlated matrix ***R***_*ν*=0.5_ + **U**′**U**, where **U** was a vector of random numbers. Different values of *p* indicate the number of biomarkers that were combined. As Table 1 clear demonstrates, both TQ and its approximation have correct sizes under varying correlation structure conditions.

The first row of Table 2 indicates the first power simulation experiments setting. Specifically, the first row demonstrates that 80% of the simulated values of the test statistic came from the null hypothesis with mean zero vector (Pr(*H*_0_) = 0.8); the remaining statistics were simulated from 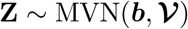, where values of mean vector were varied uniformly in the narrow interval between −0.1 and 0.1 and the correlation matrix among biomarkers **R**_*ν*_ was assumed to be equi-correlated with the common off-diagonal element *ν* = 0.5. Note, since we knew the values of **R** and b, we employed Eq. (10) to calculate 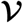 directly. The next row of Table 2 shows the number of biomarkers that were combined (i.e., 50, 30, 20, and 10), followed by the empirical power of TQ test (“Empiric. TQ”) based on evaluation of power by computing P-values under the null, and power of test based on the proposed approximation (i.e., Eq. (17), “Approx. TQ”). Since this simulation scenario matches the assumption of equi-correlated biomarkers, it is expected that our modification would result in a test at least as powerful as the original TQ-based test. The first few rows of Table 2 confirm this expectation.

The next set of simulation experiments was conducted under more realistic scenarios. Specifically, we no longer assumed that the observed biomarker levels were equi-correlated. Instead, we assumed an unstructured correlation matrix with off-diagonal elements varying between −0.25 and 0.75. To calculate power of the proposed Approx. TQ test, we substituted 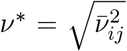 instead of ν in Eq. (17). Interestingly, as Table 2 indicates, this approximation of the general correlation structure by an equi-correlated one with the off-diagonal elements being equal to the quadratic mean of pairwise correlations did not result in appreciable drop in power. Moreover, if a small number of biomarkers were combined, our approximation resulted in higher statistical power than the traditional TQ test.

To further explore power properties, we assumed that the mean values of the test statistics under the alternative hypothesis are more heterogeneous, varying from −0.2 to 0.2. Once again, the two methods performed comparably. Finally, we increase heterogeneity among off-diagonal elements of the correlation matrix among biomarkers, *ν* ∈ (—0.70,0.90), but the two tests still had similar power performance.

The results presented in Table 2 have important implications. Specifically, based on Table 2 we can conclude that if a researcher wants to combined statistics over multiple biomarkers but does not have an access to the original data, he or she can use an average correlation value (possibly based on the previously published results) and obtained robust power values after employing our test.

### Exploring the relationship between inflamm-aging and muscle mass in senior horses

For real data application, we analyzed data from a pilot study conducted at the University of Kentucky.

It is well established that senior horses, as well as elderly humans, develop an age-related low-grade chronic inflammation termed inflamm-aging (Adams et al., 2008; Franceschi et al., 2000; Horohov et al., 2010). Elevated inflammatory markers, which characterize inflamm-aging (Franceschi et al., 2007), have been associated with diseases in elderly people, including sarcopenia (Singh and Newman, 2011), a condition defined by low muscle mass or quality, and strength (Cruz-Jentoft et al., 2018). In old horses, the relationship between inflamm-aging and such conditions as low muscle mass and strength is currently not well understood. Thus, the goal of this pilot study was to investigate the association between inflamm-aging biomarkers and muscle mass in senior horses.

### Animals

Twelve senior horses (mean = 22 years, range = 15-30 years) of light breed type and mixed sex were enrolled in this study. The horses were housed at the University of Kentucky Woodford County Farm in Versailles, Kentucky, and all animal procedures were approved by the University of Kentucky Institutional Animal Care and Use Committee.

### Muscle mass assessment

Muscle mass was determined using a previously designed, subjective, 5-point muscle development scoring system (5=best muscle development) (Walker et al., 2016). Muscle mass was evaluated by three individuals, on the left side of the horse, and scores were recorded for 6 regions: neck, thorax, abdomen, hindlimb, pelvis and lumbosacral region. In addition, body fat was scored by the same persons using a 9-point body condition scoring system (1=emaciated, 9=very fat) (Henneke et al., 1983), as fat was considered an important covariate to be included into statistical analysis related to the muscle scores.

### Inflamm-aging markers

To determine the horses inflamm-aging status, pro-and anti-inflammatory cytokines were quantified at the mRNA and protein level, in peripheral blood and in peripheral blood mononuclear cells (PBMCs), respectively. Cytokine mRNA and protein levels have been previously used to demonstrate inflamm-aging in horses (Adams et al., 2008).

For studies of cytokine mRNA, peripheral blood was collected into Tempus^™^ blood RNA tubes (Applied Biosystems, Foster City, CA) in accordance with the manufacturer’s instructions. The mRNA was extracted with an iPrep Total RNA Kit (Invitrogen, Carlsbad, CA) in an iPrep^TM^ RNA purification machine (Invitrogen), following the manufacturer’s instructions, except for resuspending the pelleted mRNA in 600*μ*l viral lysis buffer (Invitrogen). The extracted mRNA was reverse transcribed into cDNA and gene expression was quantified in the cDNA samples using real-time quantitative polymerase chain reaction (RT-qPCR), following the protocols described in Siard (2017). RT-qPCR was conducted with commercially available TaqMan assays (Thermo Fischer, Scientific, Waltham, MA) for the anti-inflammatory cytokine IL10 (Ec03468647_m1) and the pro-inflammatory cytokines IL1*β* (Ec04260298_s1), IL6 (Ec03468678_m1), IFNγ (Ec03468606_m1), TNF*α* (Ec03467871_m1), and for *β*-glucuronidase (*β*-GUS, Ec03470630_m1), a previously described reference gene (Breathnach et al., 2006). Samples were processed in duplicates and relative gene expression (RE) was calculated using the 2^−Δ*CT*^ method, where Δ*C_T_*=(Average gene of interest *C_T_* - Average reference gene *C_T_*) (Schmittgen and Livak, 2008). The RE values were used for statistical analysis.

For studies of cytokine production by PBMCs, blood was collected into heparinized blood tubes and PBMCs were isolated, counted and resuspended as previously described in Adams et al. (2008). The PBMCs were then plated in duplicates into 24 well plates at a density of 4 × 10^6^ cells ml^−1^ per well, and one well per horse received 25ng/ml of phorbol 12-myristate 13-acetate (PMA, Sigma-Aldrich, St. Louis, MO) and 1*μ*M ionomycin (IONO, Sigma) for stimulation of cytokine production. PBMCs in both wells received 10*μ*g of brefeldin A (Sigma) to stop release of protein from the golgi apparatus and thereby induce intracellular accumulation of cytokines, as required for intracellular staining. The plates were incubated for 4h at 5% CO_2_ and 37°C, and subsequently 2×200*μ*l of every well were plated into 96 well plates for intracellular staining of the pro-inflammatory cytokines TNF*α* and IFNγ, using the protocol and antibodies described in Adams et al. (2008). Once stained, the PBMCs were resuspended in 200*μ*l PBS (HyClone Laboratories, Logan, UT) and analyzed on an Attune^™^ NxT flow cytometer (Invitrogen). Lymphocytes were identified and gated based on forward scatter (FSC) and side scatter (SSC) and the percentage of TNF*α* and IFNγ positive lymphocytes after PMA/IONO stimulation was determined by selecting the fluorescence intensity of 1±0.1% positively stained lymphocytes in the unstimulated wells for each horse and recording the corresponding percent TNF*α* or IFNγ positive lymphocytes in the stimulated wells. The percentage of TNF*α* and IFNγ positive lymphocytes recorded in the unstimulated wells was then subtracted from the percentages recorded in the corresponding stimulated wells. This delta value was used for statistical analysis.

### Statistical Analysis

The initial discovery analyses focused on individual association between inflamm-aging markers and muscle scores. To explore this question, we conducted 6 regression analyses with primary outcomes defined by muscle scores at the 6 regions of the horse. To account for correlation among muscle scores induced by three individuals, we fitted linear mixed-effect models with inflamm-aging markers as main predictors, age, sex, and body condition as covariates, and evaluator as a random effect.

Next, we employed the TQ test and the approximate TQ test to combine inflamm-aging statistics across 6 body locations. To determine the appropriate correlation structure among inflamm-aging-level test statistics, we performed an additional set of simulation experiments. These simulations were important because our models at the discovery stage were slightly different from the ones presented in the Methods section. That is, instead of having multiple models with the same outcome (e.g., trait) and multiple distinct but correlated predictors (i.e., biomarkers), the current analyses involved the same set of predictors but 6 different distinct and correlated outcomes. In those simulations (not currently presented) we first confirmed that the correlation among outcomes matched the correlation among statistics obtained by linear mixed-effects models. Second, we performed an additional check of the type I error rates of the TQ and the approximate TQ tests under relatively small sample size observed in this study (*n* = 12 horses). Both the TQ and the approximate TQ tests had satisfactory test sizes (0.041 and 0.043 for the TQ and the approximate TQ, respectively). All statistical analyses were conducted using R statistical software; statistical significance was set to *P* = 0.05.

### Results

Table 3 presents results from the discovery stage. Columns labeled “%TNF*α*^+^ Lymphocytes” and “%IFNγ^+^ Lymphocytes” represent the percent TNF*α* and IFNγ positive lymphocytes after PMA/IONO stimulation and corrected for the percent recorded in the unstimulated wells. The rest of the columns represent the relative expression of pro-inflammatory cytokines (TNF*α* RE, IFNγ RE, IL6 RE, IL1*β* RE) and the anti-inflammatory cytokine (IL10 RE) in whole peripheral blood samples. The results indicate that neck scores were negatively associated with the relative expression of IFNγ (*β* = −840.818, *P* = 0.003) and TNF*α* (*β* = −298.039, *P* = 0.023) and positively with the relative expression of IL6 (*β* = 1041.320, *P* = 0.017) and IL10 (*β* = 905.628, *P* = 0.006). Lumbosacral scores were positively related to the percentage of TNF*α* positive lymphocytes (*β* = 0.018, *P* = 0.037). Hindlimb scores were negatively associated with the percent IFNγ positive lymphocytes (*β* = −0.042, *P* = 0.008) and the relative expression of TNF*α* (*β* = −268.794, *P* = 0.022), and positively with the relative expression of IL10 (*β* = 570.252, *P* = 0.045). Abdomen scores were positively associated with the percent IFNγ positive lymphocytes (*β* = 0.040, *P* = 0.044) and the relative expression of IL1*β* (*β* = 8.957 *P* = 0.013). Pelvis and thorax scores were not significantly associated with any inflamm-aging biomarker.

**Table 3:** Inflamm-aging statistics across 6 body locations obtained at the discovery stage. Individual-level associations indicate that neck scores were negatively associated to the TNF*α*, and the IFNγ RE values, and positively to the IL6 and IL10 RE values; lumbosacral scores were positively related to to the percent TNF*α* positive lymphocytes; hindlimb scores were negatively associated to the percent IFNγ positive lymphocytes and the TNF*α* and IFNγ RE values, and positively to the IL10 RE values. Abdomen scores were positively associated with the percent IFNγ positive lymphocytes and the IL1*β* RE values.

| Location | Inflammatory Markers |  |  |  |  |  |  |  |  |  |  |  |  |  |
| --- | --- | --- | --- | --- | --- | --- | --- | --- | --- | --- | --- | --- | --- | --- |
| | %TNFα<sup>+</sup> Lymphocytes | | %IFNγ<sup>+</sup> Lymphocytes | | TNFα RE | | IFNγ RE | | IL6 RE | | IL10 RE | | IL1β RE | |
| | $\beta$ | P-value | $\beta$ | P-value | $\beta$ | P-value | $\beta$ | P-value | $\beta$ | P-value | $\beta$ | P-value | $\beta$ | P-value |
| Neck | -0.014 | 0.078 | -0.03 | 0.078 | -298.039 | 0.023 | -840.818 | 0.003 | 1041.32 | 0.017 | 905.628 | 0.006 | 3.637 | 0.232 |
| Lumbosacral | 0.018 | 0.037 | -0.021 | 0.228 | 6.144 | 0.961 | -86.030 | 0.746 | 296.655 | 0.478 | 39.376 | 0.900 | -1.679 | 0.581 |
| Hindlimb | 0.005 | 0.502 | -0.042 | 0.008 | -268.794 | 0.022 | -441.42 | 0.066 | 433.769 | 0.241 | 570.252 | 0.045 | 2.448 | 0.362 |
| Abdomen | -0.003 | 0.741 | 0.040 | 0.044 | -41.405 | 0.774 | 53.068 | 0.859 | 534.718 | 0.256 | -146.099 | 0.680 | 8.957 | 0.013 |
| Pelvis | 0.014 | 0.059 | -0.023 | 0.136 | 6.941 | 0.951 | 102.851 | 0.662 | -208.343 | 0.570 | -87.896 | 0.752 | -2.874 | 0.285 |
| Thorax | 0.003 | 0.717 | -0.015 | 0.361 | -214.396 | 0.076 | -380.295 | 0.130 | 609.627 | 0.124 | 463.021 | 0.117 | 3.386 | 0.237 |

To check whether inflamm-aging markers are associated with horse’s muscle mass regardless of body location, we employed the TQ and the approximate TQ tests to combine summary statistics obtained at the discovery stage. Table 4 presents the results and indicates that only the percent IFNγ positive lymphocytes is associated with muscle condition regardless of body location. This association is likely inverse, since the individual linear mixed-effects models yielded a negative association between the percent IFNγ positive lymphocytes and muscle mass at five out of six locations (Table 3).

**Table 4:** Overall association results. Both the TQ and the Approx. TQ tests suggest that only the percentage of IFNγ positive lymphocytes is associated with overall muscle condition, regardless of body location.

| Marker | TQ | Approx. TQ |
| --- | --- | --- |
| %TNF$\alpha^+$ Lymphocytes | 0.1278 | 0.1226 |
| %IFN$\gamma^+$ Lymphocytes | 0.0435 | 0.0435 |
| TNFα RE | 0.0938 | 0.0912 |
| IFNγ RE | 0.0784 | 0.0767 |
| IL6 RE | 0.1311 | 0.1257 |
| IL10 RE | 0.0846 | 0.0826 |
| IL1 RE/$\beta$ | 0.1373 | 0.1313 |

## Discussion

In this article, we studied the problem of combining multiple correlated summary statistics. This problem has recently become increasingly important in multiple areas, including problems of pleiotropy in complex traits (Solovieff et al., 2013; Zhu et al., 2015), gene and pathway scores from SNP-phenotype association summary statistics (Lamparter et al., 2016; Liu et al., 2019), prediction of individual’s diagnostic outcome based on combination of multiple biomarkers (Mamtani et al., 2006; Yan et al., 2018), etc. We studied analytical properties of the traditional approach for combining summary statistics, TQ, which includes popular methods for association analysis of genetic variants with a disease (Pasaniuc and Price, 2017), and derived the distribution of the TQ-statistic under the alternative hypothesis. Based on our derivation, we proposed a scale-location approximation of the TQ statistic, which we called approximate TQ. We showed that the approximate TQ has very similar power to the traditional TQ test under varying correlation structures among biomarkers. This finding is not trivial and has practical implications. For example, much of the interest in gene-based tests has recently centered on the simulation-based methods such as VEGAS (Liu et al., 2010). Without access to individual-level data, simulation-based methods incorporate LD information from reference panels (e.g., HamPMap2 (Consortium et al., 2007) or 1,000 Genomes data (Consortium et al., 2012)) to account for the correlation among statistics. However, sensitivity of these methods to misspecification of the LD pattern were poorly understood. Our results suggest that the use of reference panels in the absence of genotype data or even just a single value of quadratic mean of pairwise correlations results in adequate power and type I error rates of the combination test.

Based on previous studies investigating the relationship between inflammatory markers and muscle mass in other species (Toth et al., 2006; Visser et al., 2002; Wang et al., 2014), we expected to observe a positive association between the anti-inflammatory marker (IL10 RE) and muscle mass, and a negative association between the pro-inflammatory markers (%IFNγ^+^ and %TNF*α*^+^ lymphocytes, and IL6 RE, IL1*β* RE, TNF*α* RE and IFNγ RE) and muscle mass. The initial discovery analysis showed, however, that pro-inflammatory markers were both positively and negatively associated to muscle scores, and that the biomarker associations were not consistent across all scoring locations. This observation might illustrate the complex relationship between muscle mass and inflamm-aging markers, and suggest that this relationship could be indirect or non-causative; especially for blood-level inflamm-aging markers, which might be less intimately linked to muscle mass than inflammatory makers in the muscle (Wilson et al., 2017). n addition, the mRNA and in vitro study-based inflamm-aging markers used in this pilot work might be less closely related to muscle mass than circulating cytokine proteins, which have been found associated to muscle mass in previous studies (Toth et al., 2006; Visser et al., 2002). The observed associations are furthermore complicated by the fact that IL6, considered a pro-inflammatory cytokine in this study, can have both pro-and anti-inflammatory effects, depending on the receptor it engages (Scheller et al., 2011).

Results from TQ and approximate TQ analysis showed a significant, and likely inverse, association between the percent IFNγ positive lymphocytes and muscle mass across all scoring locations. This finding does not corroborate the results of a previous study investigating the relationship of muscle mass and inflamm-aging in senior horses. In this previous study the percent IFNγ positive lymphocytes was tested for correlation with overall muscle mass as well, and no significant association was found (Siard, 2017). These contradicting results might originate from different muscle mass assessment methods used. While we applied a muscle development scoring system leading to six regional scores (Walker et al., 2016), Siard (2017) used a picture based whole body muscle scoring system (Graham-Thiers and Kronfeld, 2005), and ultrasound and deuterium oxide dilution based fat free mass estimations (Dugdale et al., 2011; Lehnhard et al., 2004). Furthermore, different outcomes of these two studies may be due to analysis of the data using different statistical methods, as well as using different age-range of horses studied (range: 15-30 yrs in the current pilot study, vs range: 18-29 yrs in Siard (2017)).

To our knowledge this is the first study suggesting a potential relationship between inflamm-aging and muscle mass in senior horses. However, given the contradictory findings in another study, and the ambiguous association of markers depending on the scoring location, further investigation is imperative. We also note that the traditional statistical approaches (e.g., linear regression and linear mixed-effects models) could only explore the relationship between inflamm-aging and muscle mass at one assessment location at a time, due to the relatively small sample size (i.e., one would run out of the degrees-of-freedom if were to incorporate all muscle mass assessment locations into a single model). The combination test provided an elegant solution to this statistical issue. We stress that our application is one of many data sets typically encountered in modern research, when the number of biomarkers (usually several thousands) greatly exceeds the number of samples (usually in hundreds or less) and a single full model can not be built. Furthermore, the TQ and the approximate TQ are generally applicable to situations, in which correlated summary statistics need to be combined into an overall score. The power of individual summary statistics can often be weak but they can re-enforce the effect of one another in a combination test, improving the overall statistical power.

## Author contributions

DZ and OV performed theoretical derivations; AH, BC, CB, DB and AA conducted the pilot study and DZ, OV, AH and AA drafted the manuscript.

